# Increased sensorimotor activity during categorisation of ambiguous emotional expressions

**DOI:** 10.1101/717553

**Authors:** Ozge Karakale, Matthew R. Moore, Nicolas McNair, Ian J. Kirk

**Affiliations:** School of Psychology, The University of Auckland, Auckland, New Zealand; School of Medicine, The University of Auckland, Auckland, New Zealand; School of Psychology, The University of Sydney, Sydney, Australia; Centre for Brain Research, The University of Auckland, New Zealand; Brain Research New Zealand

**Author notes:** Corresponding author: Ozge Karakale, School of Psychology, The University of Auckland, Private Bag 92019, Auckland 1142, New Zealand. E-mail address (O. Karakale).

**Keywords:** EEG, mu suppression, mirror neuron, sensorimotor activity, facial expression, predictive coding

## Abstract

Actions are rarely devoid of emotional content. Thus, a more complete picture of the neural mechanisms underlying mental simulation of observed actions requires more research using emotion information. The present study used electroencephalography (EEG) to investigate mental simulation associated with facial emotion categorisation. Mu rhythm modulation was measured to infer the degree of sensorimotor simulation. Categorising static images of neutral faces as *happy* or *sad* was associated with stronger mu suppression than categorising clearly happy or sad faces. Results suggest the sensitivity of the sensorimotor activity to emotional information rather than visual attentional engagement, because further control analyses revealed (1) no effect of emotion type on occipital alpha modulation, and (2) no difference in mu modulation between the conditions of a control task, which required categorising images with the head oriented right, left, or forward as *right* or *left*. This finding provides evidence for the role of the sensorimotor activity in a higher form of mental simulation. Consistent with a predictive coding account of action perception, stronger mu suppression during attempted emotion categorisation of neutral faces may involve minimising the mismatch between predicted kinematics of a happy/sad face and the observed stationarity of neutral faces.

## 1 Introduction

Mental simulation theories of action understanding suggest that observed motor acts, defined as a series of goal-related movements, are matched onto the motor system of the observer, thereby allowing an understanding of the intentions behind actions (Gallese et al., 2009). Simulation, however, describes the mapping of not only movements, but also emotions and sensations of others onto the observer’s visceral and somatosensory systems (Gallese & Sinigaglia, 2011). It is this process of external-to-internal mapping that forms the basis of understanding others, including recognising motor actions and inferring their goals, as well as understanding internal states and motivations (Gallese & Sinigaglia, 2011). Successful operationalisation of this mapping process involves reducing any ambiguity present in an observed action to achieve a correct understanding of the intent behind the act being performed. In the context of action perception, ambiguous information can be described as movements that are open to multiple interpretations. Myriads of actions and emotions exhibited in social environments make the skill to resolve ambiguity of social emotion information crucial to understanding others.

Mental simulation that is specifically associated with the activity of the motor system during action observation is called *motor simulation* or *motor resonance* (Gordon et al., 2018). A possible biological basis of motor simulation has been proposed with the discovery of a special category of neurons, called *mirror neurons*, that are active during both action execution and observation in monkeys (di Pellegrino et al., 1992; Dushanova & Donoghue, 2010; Gallese et al., 1996; Rizzolatti et al., 1996). These motor neurons, mostly selective for a particular movement configuration, such as grasping an object or breaking it, were observed to discharge not only while the monkey executed certain actions, but also while they observed the experimenter perform the same or similar ones (Gallese et al., 1996; Rizzolatti et al., 1996). Due to this response characteristic of mirror neurons, it was argued that they could provide a mechanism for coding the internal motor representations of actions in the motor system which are triggered by perception as well as execution of actions. Through the strengthened connections formed as a result of repeated pairings between the internal motor representations and their consequences, representations of an action activated during its observation would enable access to the knowledge of that action’s consequences. Based on this rationale, it was argued that mirror neurons allow understanding actions and their goals (Gallese et al., 1996; Rizzolatti et al., 2001). These types of neurons have also been discovered in humans (Mukamel et al., 2010), and neuroimaging studies have demonstrated that similar patterns of activation in the human brain are generated during action execution and observation (Molenberghs et al., 2012).

Motor simulation in humans has also been investigated using electroencephalography (EEG) and magnetoencephalography (MEG). These studies have identified oscillatory activity in the alpha (8 – 13 Hz) and beta (14 – 25 Hz) frequency bands, originating in motor and somatosensory cortices, as potential neurophysiological markers of action understanding (e.g., Caetano et al., 2007; Simon & Mukamel, 2016; Ulloa & Pineda, 2007). The alpha frequency activity generated by the sensorimotor cortex, also called the *mu* rhythm, is most prominent when the motor system is at rest, and is suppressed by the execution, observation, or imagery of limb and face movements (Cochin et al., 1998; Cochin et al., 1999; Debnath et al., 2019; Lepage & Theoret, 2006; Muthukumaraswamy et al., 2004; Muthukumaraswamy et al., 2006; Sakihara & Inagaki, 2015). These findings support the existence of a mirror-neuron network (MNN) in humans that is implicated in action perception (for a recent review, see Keysers et al., 2018).

In addition to motor simulation, evidence for a similar mechanism underlying emotion processing has arisen from research showing an overlap between neural structures that are activated during voluntary imitation and observation of emotional facial expressions; specifically, the premotor area, the parietal area, the insula and the amygdala (Carr et al., 2003; Leslie et al., 2004; Van der Gaag et al., 2007). Consistent with a motor theory of empathy, these findings indicate that action representations that are activated during facial expression observation communicate with the limbic areas through the insula, possibly allowing for understanding and sharing emotions (Carr et al., 2003). This concordance between perception and sensation has been observed in a variety of sensory domains, including pain (Corradi-Dell’Acqua et al., 2011; Hutchison et al., 1999; Singer et al., 2004), touch (Keysers et al., 2004; Kuehn et al., 2014; Lamm et al., 2015), disgust (Calder et al., 2000; Wicker et al., 2003), and taste (Jabbi et al., 2007). Taken together, findings indicate that mirroring is very likely to also be a property of non-motor systems in various cortical and subcortical brain regions.

Mu rhythm activity has been found to be sensitive to top-down contextual influences. Studies that have investigated the modulatory effect of pain on the mu rhythm have reported greater suppression during visual perception of limbs in painful compared to non-painful situations (Cheng et al., 2014; Hoenen et al., 2015; Yang et al., 2009). For example, recently, Hoenen et al. (2018) demonstrated that viewing actions (e.g., cutting a cucumber) depicting pain (e.g., finger placed between knife and cucumber) evoked stronger mu suppression than their almost visually identical no-pain counterparts (e.g., finger not endangered). Other studies have found that the relevance of the observed action for the observer modulates motor resonance (Dickter et al., 2013; Oberman et al., 2008; Varnum et al., 2016). For example, Dickter et al. (2013) found greater mu suppression in smokers than non-smokers to viewing humans interacting with cigarettes than only cigarette images, whereas both groups showed greater mu suppression while viewing nonsmoking-related stimuli (e.g., toothbrush) in the interaction than the object-only condition. In children, viewing hand movements showed the greatest mu suppression to their own movement, followed by a familiar person’s, and least suppression to the stranger’s, indicating the sensitivity of the mu rhythm to the actor’s familiarity (Oberman et al., 2008). In a study investigating the link between mu suppression, reward value of the stimuli and empathy, Gros et al. (2015) reported stronger mu suppression in the alpha and beta bands while viewing happy expressions that had been conditioned to be associated with a reward compared to those associated with a loss. This finding suggests that empathy towards the actor modulates mu suppression. The sensitivity of the mu rhythm to contextual information indicates that the internal simulation of actions extends beyond crude motor mimicry, which involves merely mirroring the kinematic properties of observed actions and suggests an assessment of the social content of the stimuli.

Recent contributions to our understanding of emotion simulation have come from work exploring mu modulation during observation of different types of facial expressions. In a study in infants, Rayson et al. (2016) found significant bilateral mu suppression for non-emotional expressions (mouth opening), but only right-hemispheric suppression for emotional (happy or sad) facial expressions. An emotion categorisation study from our group reported greater mu suppression while viewing dynamic videos of non-emotional facial movement (mouth opening) compared to non-biological movement (kaleidoscope pattern), but this was not observed for emotional faces (happy or sad) (Karakale et al., 2019). Greater mu suppression in the mouth opening condition may be due to the heavier workload imposed on the sensorimotor system when attempting to extract emotional content from a non-emotional stimulus. This contrasts to the relative ease in inferring happiness/sadness from the very pronounced happy/sad facial expressions. This suggests that the mental simulation system is engaged more by ambiguous social information compared to readily recognisable actions and emotions (Karakale et al., 2019). Several infant studies have also demonstrated that watching actions that do not have readily recognisable meanings (e.g., bringing a cup to the ear, turning on a lamp with head) evoked stronger mu suppression than ordinary actions (bringing a cup to the mouth, turning on a lamp with hand) (Langeloh et al., 2018; Stapel et al., 2010). These findings indicate that the sensorimotor system is involved in deciphering the meaning of actions. The sensitivity of the mu rhythm to ambiguous social emotion information is a question that remains to be resolved and is key to understanding the role of the sensorimotor system in emotion processing.

The role of the sensorimotor cortex in emotion understanding can be modelled through the *Predictive Coding Framework* of action perception (Kilner et al., 2007; Kilner, 2011). This model conceives of actions at four different levels of abstraction: *intention* (the long-term goal initiating the action), *goals* (the short-term objective of the component action), *kinematics* (the sensory input providing information about the trajectory and the velocity of the action), and the *muscle level* (the muscles directly used to execute the action). At each level, the *predicted attribute*, generated by past experiences and prior expectations, gets compared to the *observed attribute*, which is defined by the immediate incoming information. For example, *predictions* about the objective of an action are sent from the goal level down to the kinematics level, and there the predicted kinematics associated with that goal get compared to the observed kinematics. Any discrepancy between a prediction and an observation generates a *prediction error*. This error signal is propagated back to higher levels, and results in *prediction updating* in the goals or even intentions of the action. Via continuous reciprocal interactions between the different levels, the prediction error is minimised and a more accurate understanding of an observed action is determined. Through this predictive mechanism, the observer is able to tell, for example, that the observed individual is picking up a cup in order to drink from it. However, if after picking up the cup the person tips the contents into a sink, prediction error would need to be minimised through updating of goals and intentions to arrive at a new understanding of the action (e.g., to clean the cup).

A similar predictive process may be engaged when deciphering the emotional content of an action. Based on their knowledge and experience about the mental states associated with different types of smiles, an observer might interpret a smile in a face which involves lip pressing as affiliative, therefore arrive at an understanding of the action intention as signaling an interest in maintaining a social bond (posterior belief). However, if the observer then sees a subtle pursing of the lips contradicting their initial interpretation, this sensory information would be used to update their belief through prediction error minimisation. At the kinematics level, this would involve reducing the discrepancy between the new sensory information about the kinematics of the observed movement and the prediction sent from the higher levels signaling the kinematics of an affiliative smile. When the error is minimised at all levels of the hierarchy through the reciprocal connections, this particular action of smiling could finally be interpreted as signaling dominance. From this perspective, expected, predicted or easily recognisable emotional expressions would be associated with a smaller prediction error, whereas others would cause a larger error. Larger prediction errors would require stronger employment of the mental simulation mechanism to reduce the level of uncertainty about this information.

The aim of this study was to investigate the effect of ambiguity in social affective stimuli on mental simulation. For this purpose, we measured the modulation of sensorimotor mu activity during categorisation of happy, sad and neutral facial expressions. As described previously, past research shows that mu modulation likely reflects a higher-order simulation process rather than a simple mirroring of movement kinematics. Thus, although faces with clear, pronounced emotional expressions (‘unambiguous’) possess greater implied facial movement than those with more subdued, subtle expressions (‘ambiguous’), we in fact predict that there would be greater mu suppression in response to the ambiguous expressions. This is because the ambiguity inherent to neutral expressions leads to a greater workload being imposed on the sensorimotor system as it attempts to simulate the emotional content. To establish that any effects were specific to evaluating the emotional content of the faces, we included a control task that required participants to judge the direction in which faces were looking. Since this does not require mental simulation, the direction the faces are looking should not affect mu suppression. Finally, in order to ensure our data specifically reflected mu activity from the sensorimotor cortex, we also measured alpha activity over the occipital cortex. This posteriorly-generated alpha is modulated by a number of non-motor factors, such as visual processing and attention (Hobson & Bishop, 2016), but should not differentiate between faces with differing expressions or viewpoints.

## 2 Methods

### 2.1 Participants

Twenty-three neurologically-intact participants (12 female, mean age = 24, *SD* = 4.9 years), with normal or corrected-to-normal vision, were recruited through advertisements placed around the University of Auckland campus and online through the School of Psychology website. Participants were compensated for their time with a $20 supermarket voucher. All participants provided written informed consent, and the experimental protocol was approved by the University of Auckland Human Participants Ethics Committee and conducted in compliance with the Declaration of Helsinki.

### 2.2 Stimuli

Stimuli were obtained from the Radboud Faces Database (RaFD; Langner et al., 2010). The stimulus set contained static facial images of 38 actors (19 female) portraying *happy, sad* and *neutral* facial expressions for the Emotion task, and *left-, right-* and *forward-oriented* faces for the Orientation task. Each image was presented only once in each condition. Images were converted to greyscale and their size corresponded to 7.9° × 8.2° cm visual angle at a viewing distance of about 57 cm.

### 2.3 Experimental tasks and procedure

In the Emotion task, participants categorised *happy, sad*, and *neutral* forward-oriented faces as either *happy* or *sad*. In the Orientation task, they categorised *left-oriented* (turned 45° left), *right-oriented* (turned 45° right), and *forward-oriented* neutral face images as either *left* or *right*. Both tasks used an identical protocol (see *Figure 1*). Each trial started with a 2000 ms blank screen, before a fixation cross appeared at the centre of the screen. The duration of this fixation varied randomly between 1050 and 1350 ms. Following the fixation cross, a face stimulus was presented at the same location for 1000 ms. Then, a blank screen appeared for 1000 ms, followed by the response slide. This slide presented the two possible responses on the left and right of the screen: *happy-sad* in the Emotion task; or *left-right facing arrows* in the Orientation task. Participants pressed the “d” key on a keyboard if the emotional expression (Emotion task) or head orientation (Orientation task) more closely matched the label on the left, and the “k” key if it more closely matched the label on the right. The response prompt remained on the screen until a response was made before progressing to the next trial. The two tasks were completed in separate blocks, with the order counterbalanced across participants. Each task contained 38 images in each of their 3 conditions (*happy/sad/neutral* or *left/right/forward*), for a total of 114 trials in each block. Participants were allowed a 5- to 10-minute rest period between tasks, and the whole experiment lasted about 30 minutes.

**Figure 1.**
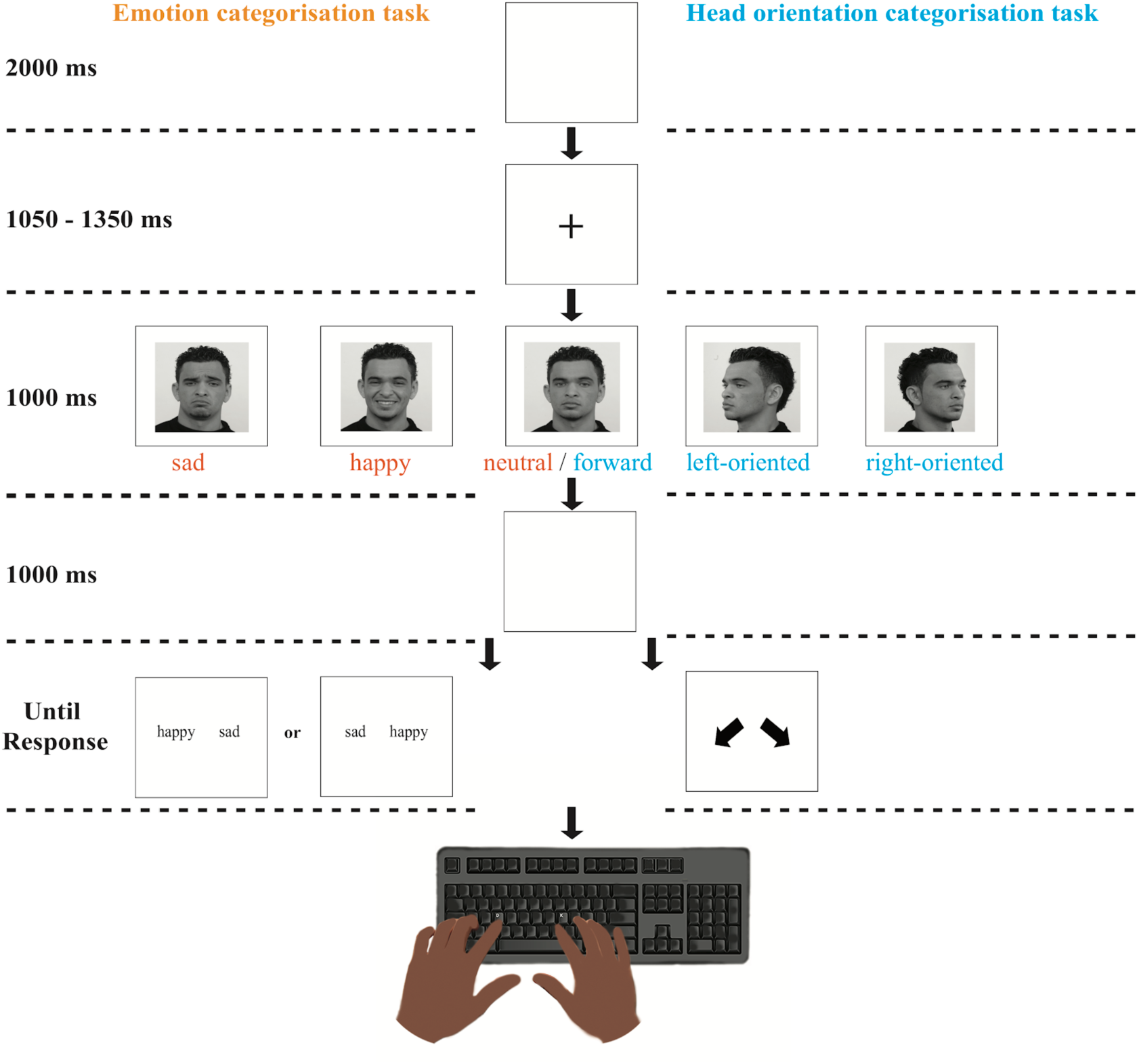
Structure of the experimental design. Shown down the centre are the slides that were common to both tasks. Example stimulus images for each of the conditions for the emotion categorisation task are presented on the left and centre (orange text), and for the head orientation categorisation task on the centre and right (blue text). Each trial started with a blank screen (2000 ms). Then a fixation cross appeared at the centre of the screen (1050 - 1350 ms), followed by a face stimulus (1000 ms). After this, a blank screen appeared (1000 ms), before participants were prompted for their response. The response slide presented the labels *happy* and *sad* in the Emotion task or *left-* and *right-facing arrows* in the Orientation task. Participants pressed “d” key on a keyboard if the emotional expression or the head orientation of the face matched the label on the left, and the “k” key if it matched the label on the right. The next trial began as soon as a response was made.

Happy and sad faces in the Emotion task, and right- and left-oriented head images in the Orientation task were designated *unambiguous* conditions. These stimuli had a greater degree of implied movement relative to the neutral forward-oriented faces, which were classified as the *ambiguous* condition and were identical between the two tasks. Given the two-alternative, forced-choice nature of the tasks, categorisation of the ambiguous stimuli was therefore completely arbitrary. However, participants were only told that some of the expressions/orientations would be easy to categorise whereas others would be more difficult. The fact that some faces contained no emotion or were oriented directly forwards was not made explicit to the participants. During debriefing, some participants expressed finding some images difficult to categorise, but none mentioned the fact that they were neutral or forward-facing.

### 2.4 EEG data recording

During the experiment, participants sat in a comfortable chair in an electrically- and acoustically-shielded, dimly-lit room (IAC Noise Lock Acoustic – Model 1375, Hampshire, United Kingdom). EEG was recorded continuously at 1000 Hz with an analogue band-pass filter of 0.1 - 400 Hz, using a 128-channel Ag/AgCl electrode net (Electrical Geodesics Inc., Eugene, Oregon, USA). Electrode impedances were kept below 40 kΩ, an acceptable level for this system (Ferree et al., 2001). EEG was acquired using a common vertex (Cz) reference and later re-referenced to an average reference offline.

### 2.5 EEG data preprocessing

EEGLAB toolbox (v13.6.5b; Delorme & Makeig, 2004) was used to preprocess the EEG data in MATLAB R2017a (Mathworks Inc.). First, data were downsampled to 250 Hz, and then high-pass filtered at 0.1 Hz and low-pass filtered at 40 Hz. The continuous recordings were then segmented into 4000 ms epochs, extending from 2000 ms before to 2000 ms after stimulus onset. Channels with an absolute threshold or activity probability limit of 5 SD, based on kurtosis, were identify as bad and replaced with an interpolation using the other channels. Data were then re-referenced to the average before performing an independent component analysis (ICA). The first 10 ICA components were inspected and any components representing eye blink or eye movement artefacts were removed.

### 2.6 EEG data analysis

For each task and for each condition, 1000 ms baseline epochs corresponding to the fixation cross immediately preceding stimulus onset, and 1000 ms stimulus epochs starting from stimulus onset were obtained. Analysis of mu (8 – 13 Hz) power was conducted using data from 12 central electrodes overlying primary sensorimotor areas (left: C3, 30, 31, 37, 41, 42 and right: C4, 80, 87, 93, 103, 105), and for alpha power using data from 3 occipital electrodes over the visual cortex (O1, Oz and O2) (see *Figure 2*). At each electrode, a Fast Fourier Transform was applied to calculate the power spectral density (PSD) associated with the baseline period and the stimulus period of each trial. Mu/alpha power modulation at each electrode was calculated by taking the ratio of the stimulus period PSD relative to that of the baseline period. Trials with a PSD ratio value greater than 3 scaled median absolute deviations (MAD) from either the central or the occipital cluster median PSD ratio value were excluded as outliers. In order to obtain the mu and alpha modulation scores for each condition, the mean of the PSD ratio values of the central and occipitals clusters was calculated, respectively. This resulted in a single mu and a single alpha ratio score for each condition, for each participant. Due to non-normal distribution of ratios, data were log-transformed prior to statistical analysis. A log-ratio value of less than zero indicates mu/alpha suppression, while a value greater than zero indicates enhancement.

**Figure 2.**
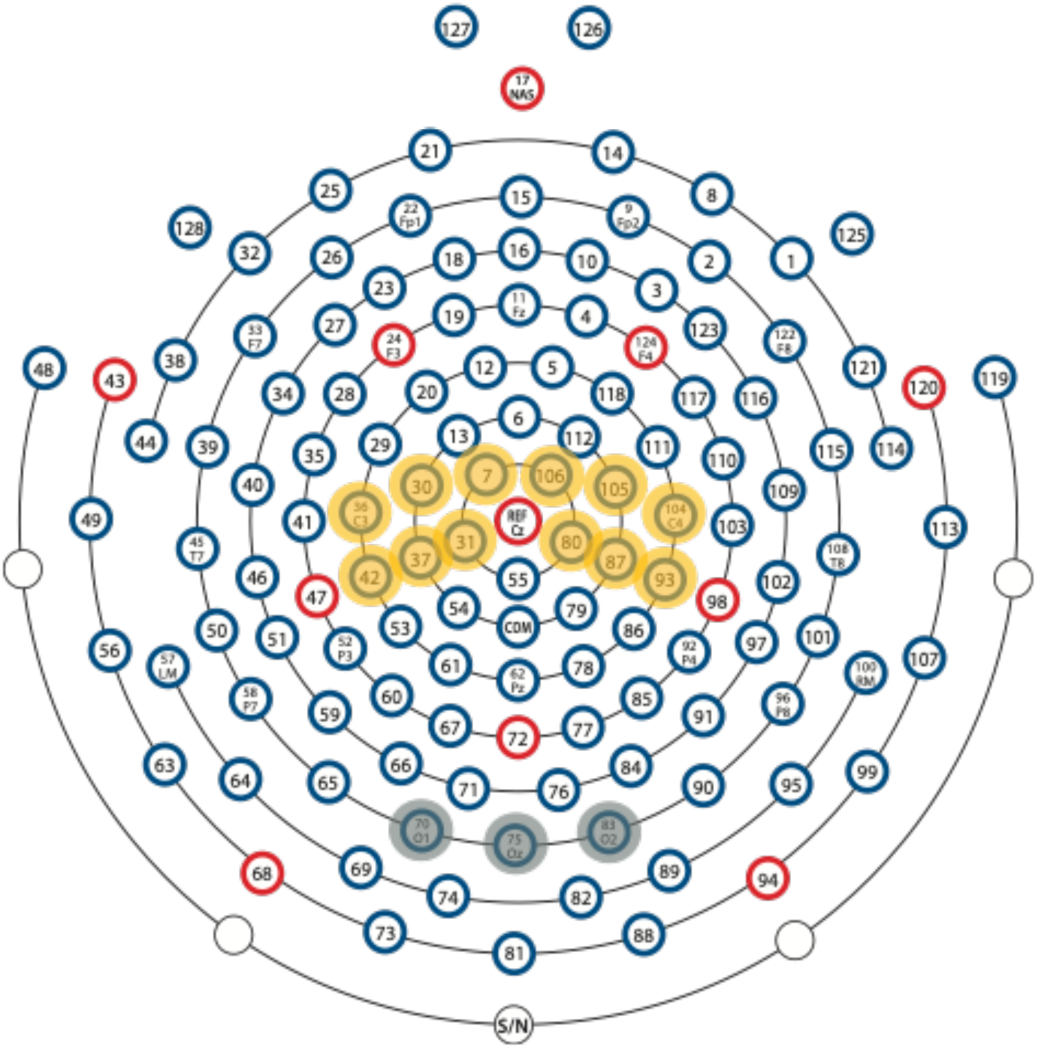
EEG electrode clusters used for statistical analysis. Top view of the 128-channel EEG array shows the central cluster used to calculate mu power modulation (highlighted in yellow), and the occipital cluster used to calculate alpha power modulation (highlighted in grey).

Upon visual inspection of preprocessed EEG data, recordings from one participant were identified as extremely noisy and so was not analysed further. In addition, during data analysis 2 participants during the Emotion task and 4 participants during the Orientation task had one or more PSD ratio values that were greater than 3 scaled MAD from the median PSD ratio value. Therefore, they were excluded from statistical analysis. Finally, accuracy in the unambiguous conditions of both tasks was calculated for each participant to ensure that they had paid attention throughout the task. One participant incorrectly categorised 16% and 26% of the trials in the left and right head orientation conditions, respectively. This differed from the other participants, for whom accuracy was close to 100%, and so they were removed from further analysis. The following statistical analyses were conducted on data from 19 participants in the Emotion task and 17 participants in the Orientation task. R (R Core Team, 2019) was used for all statistical analyses, except the exploratory event-related spectral perturbation (ERSP) analysis, which was performed using the EEGLAB toolbox (v2019.0; Delorme & Makeig, 2004) in MATLAB R2018b (Mathworks Inc.). For all tests, a *p*-value of < .05 was assumed to be statistically significant.

## 3 Results

Before comparing between task conditions, the degree of mu and alpha modulation within each condition was assessed in a series of paired-samples *t*-tests by comparing the stimulus period PSD values with their corresponding baseline period PSD values. In the Emotion task, significant mu suppression was observed in response to both *neutral* (*t*(18) = 4.88, *p* < .001) and *happy* (*t*(18) = 2.44, *p* = .025) faces, but only marginally to *sad* faces (*t*(18) = 2.07, *p* = .053). Significant occipital alpha suppression was observed to all faces (*happy*: *t*(18) = 2.42, *p* = .026; *sad*: *t*(18) = 3.17, *p* = .005; *neutral*: *t*(18) = 3.31, *p* = .004). In the Orientation task, significant mu suppression was observed to *forward*-oriented faces (*t*(16) = 4.84, *p* < .001), but not *left* (*t*(16) = 1.87, *p* = .08) or *right* (*t*(16) = 1.54, *p* = .144) faces. Significant occipital alpha suppression was observed to faces oriented *forward* (*t*(16) = 2.37, *p* = .031) and *right* (*t*(16) = 2.14, *p* = .048), and marginally to *left* (*t*(16) = 1.95, *p* = .069).

Next, four separate one-way repeated measures ANOVAs were run to assess the degree of difference in mu and alpha modulation in the Emotion and the Orientation tasks. Results from Mauchly’s test indicated that the assumption of sphericity was met in all instances. There was a significant main effect of emotional expression on mu modulation (*F*_(2, 36)_ = 3.5, *p* = .041, *η*_G_^*2*^ = 0.08), but not of head orientation (*F*_(2, 32)_ = 1.73, *p* = .193, *η*_G_^*2*^ = 0.03). Two one-tailed planned comparisons (i.e., neutral > happy, neutral > sad) then tested our hypothesis that mu suppression would be significantly greater in the ambiguous condition (i.e., *neutral*: *M* = −0.041, *SD* = 0.064) compared to either of the unambiguous conditions (i.e., *happy*: *M* = −0.005, *SD* = 0.078; or *sad*: *M* = 0.012, *SD* = 0.087). Consistent with our hypothesis, mu suppression was significantly greater in response to *neutral* faces compared to *happy* (*t*(18) = 1.85, *p* = .041) and *sad* (*t*(18) = 2.22, *p* = .02) faces. As predicted, there was no difference between *happy* and *sad* expressions (*t*(18) < 1). Furthermore, there was no effect of head orientation on mu suppression (*F*_(2, 32)_ = 1.73, *p* = .193, *η*_G_^*2*^ = 0.03), and there were no differences in the amount of occipital alpha suppression between conditions of either task (Emotion: *F*_(2, 36)_ = 1.166, *p* = 0.323, *η*_G_^*2*^ = 0.01; Orientation: *F*_(2, 32)_ = 2.18, *p* = .129, *η*_G_^*2*^ = 0.01) tasks. The distribution of the mu/alpha modulation scores are displayed in *Figure 3*.

**Figure 3.**
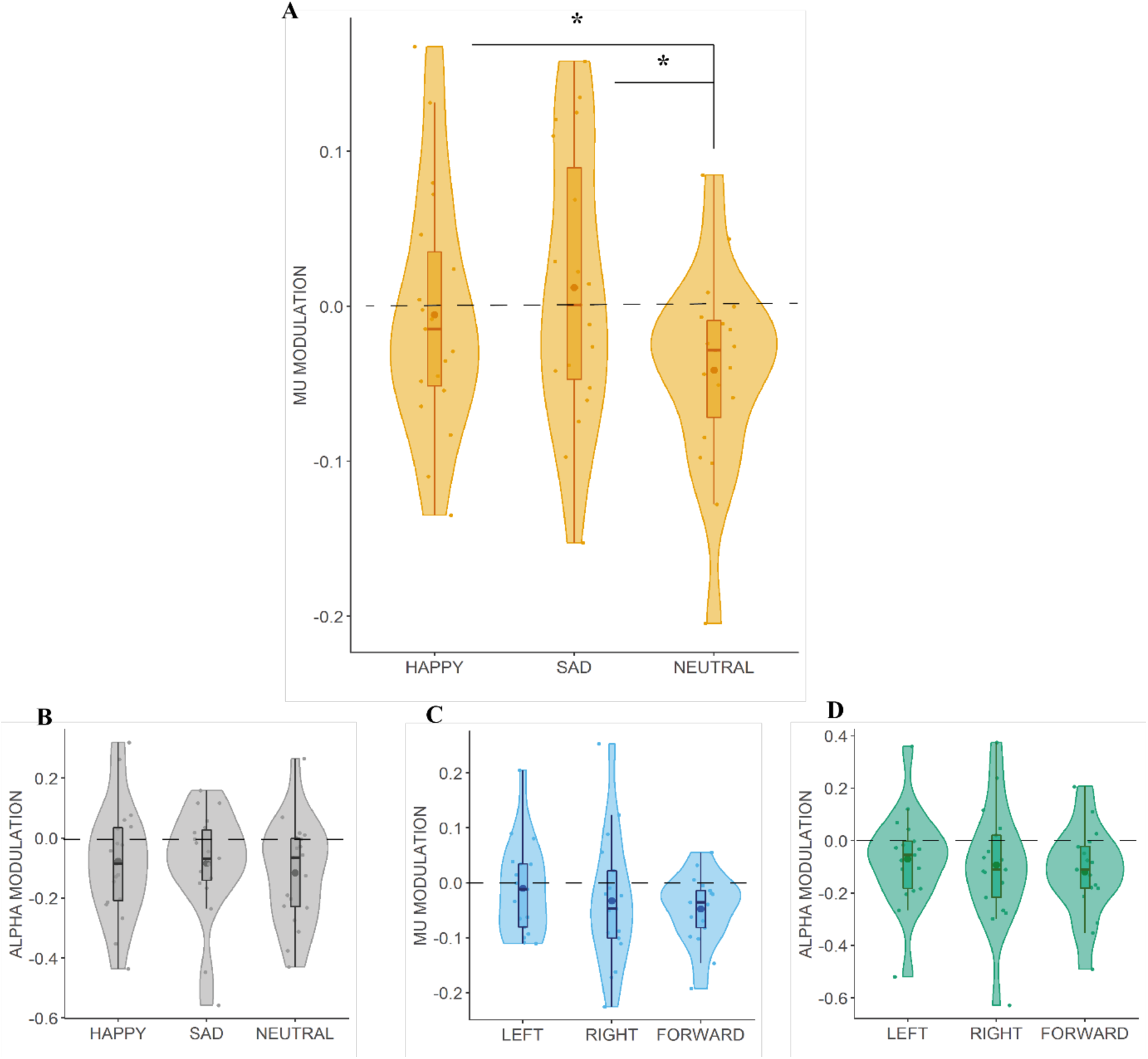
Modulation of (A) mu and (B) alpha in the emotion task, and (C) mu and (D) alpha in the head orientation task. Each dot corresponds to a participant’s log PSD ratio. Scores below 0 indicate suppression and above 0 indicate facilitation. Inside each boxplot, the bigger dot represents the mean and the horizontal line represents the median of the condition. Kernel probability density of the data at different ratio values are represented by the density plots surrounding the data points. Significant differences are marked by an asterisk (* *p* < .05).

Further exploratory analyses on ERSP in the Emotion and the Orientation tasks were conducted on four representative channels, two over the sensorimotor (C3 and C4) and two over the occipital cortex (O1 and O2). *Figure 4* illustrates the mean ERSP associated with stimulus processing during both tasks, calculated over the time window of approximately 1000 ms pre- and post-stimulus. The time-frequency points that deviated significantly from the baseline were determined using parametric statistics (*p* < .05, uncorrected). In the Emotion task, significant differences between conditions in suppression of power in the 8 - 13 Hz frequency band were observed beginning with stimulus onset (0 ms) at both sensorimotor channels. The plots suggested that this difference arose from greater suppression to *neutral* compared to *happy* and *sad* faces (top left). Other plots did not suggest any differences between conditions which appeared to be driven by greater alpha band suppression to *neutral* than *happy* and *sad* faces at O1 and O2 (bottom left), or to *forward-* compared to *left-* and *right-oriented* faces at C3 and C4 (top right) or at O1 and O2 (bottom right).

**Figure 4.**
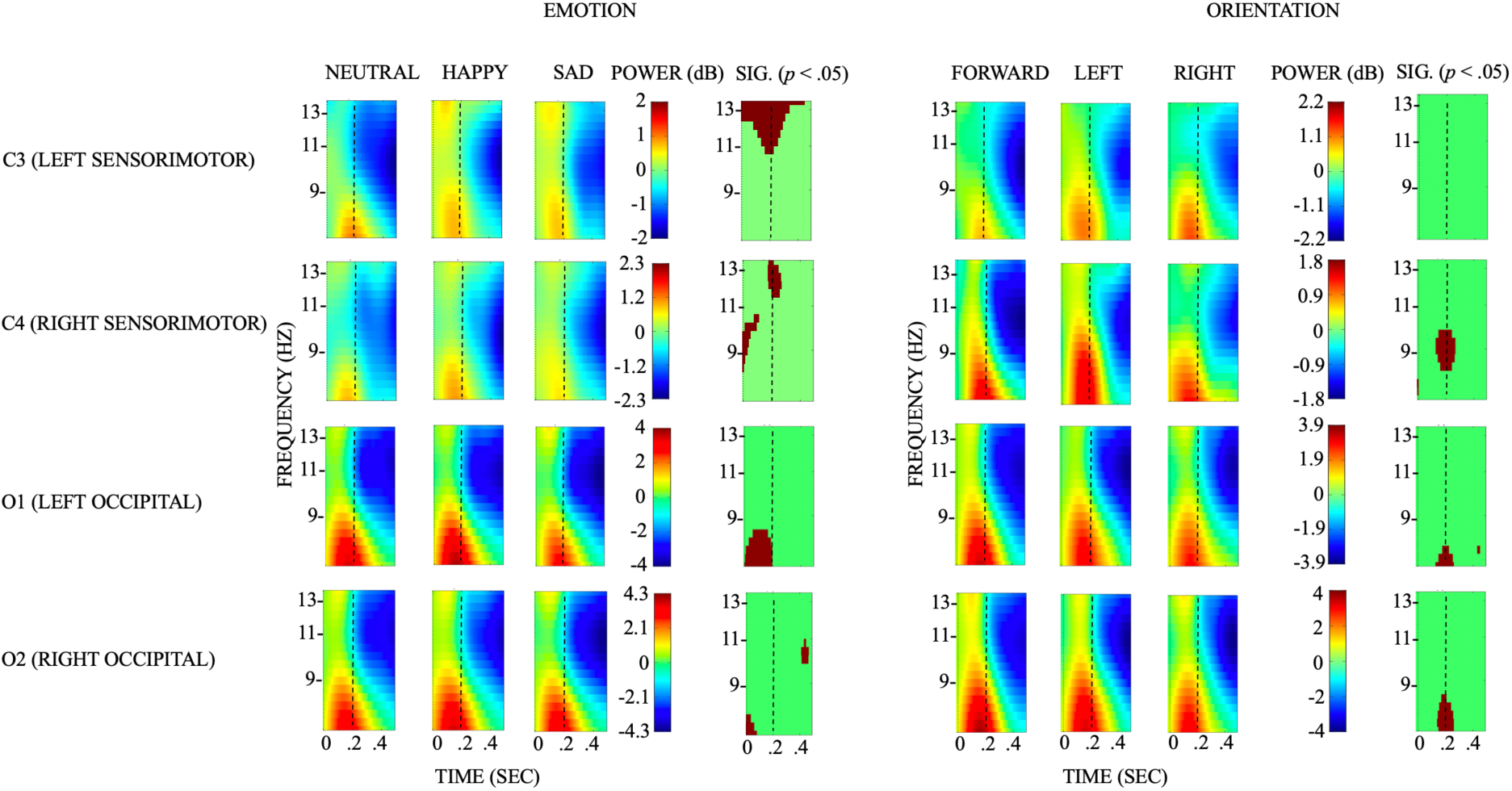
Event-related spectral perturbation (ERSP) associated with emotion or orientation categorisation relative to the 1000 ms pre-stimulus baseline. ERSP plots from representative electrodes selected over the regions of interest (i.e., left and right sensorimotor and occipital cortices) show the time-frequency decomposition for each channel (i.e., C3, C4, O1, O2) in each task. Colour bars depict power in dB. The ERSP plots depict significant differences (*p* < .05, uncorrected) between conditions in their alpha band suppression levels at channels C3 and C4 during the Emotion task, which appears to arise from greater suppression beginning with stimulus onset (0 ms) in the neutral face condition compared to happy and sad faces.

## 4 Discussion

### 4.1 General discussion

The goal of this study was to investigate the sensitivity of sensorimotor simulation to ambiguous facial expressions. To do this, we measured EEG mu suppression as participants viewed unambiguous (happy/sad) and ambiguous (neutral) facial expressions. As hypothesised, there was significantly greater mu suppression over sensorimotor regions in response to neutral faces compared to happy or sad faces. Given the link between mental simulation and mu suppression, this suggests greater involvement of the MNN when evaluating the emotional content of neutral faces. A generalised alpha suppression to visual differences between neutral and emotional faces was excluded as occipital alpha showed no difference between the different expressions. Furthermore, the mu suppression was not the result of some form of a general task effect, as determining the orientation of the faces’ viewpoints produced no similar mu suppression.

The selective response of mu oscillations to categorisation of neutral faces suggests that it is not purely responding to the amount of movement, here implied, in the stimulus. Past work has found that the mu rhythm is sensitive to factors beyond movement kinematics, such as the contextual appropriateness of motorically-identical actions (Koelewijn et al., 2008). Stronger mu suppression to neutral faces in the present study, which lacked almost any implied movement compared to happy or sad facial expressions, provides evidence for the involvement of the sensorimotor cortex in higher-level cognitive processing of emotion. Assessing the emotional content of neutral faces is not as easily determined as it is for purely happy and sad faces. As such, sensorimotor processing might become more intense as it attempts to resolve this ambiguity. These results suggest that mental simulation of emotion is more taxing when processing faces with expressions that are not easily mapped to a particular emotion.

Observing ambiguous social information, as in the current study and Karakale et al. (2019), incorrect execution of actions (Koelewijn et al., 2008), or unusual actions, such as turning a lamp with the head (Langeloh et al., 2018) or lifting a cup to the ear (Stapel et al., 2010), appears to increase activity in implicated brain networks to resolve the issue of violated prior action expectations. In the predictive coding framework, action predictions that are violated by incoming sensory information are updated through error minimisation, which is a process reflected in enhanced activity in the motor system (Gardner et al., 2015; Gerson et al., 2017; Gordon et al., 2018; Kilner et al., 2007). If the discrepancy between the prediction at one level and the subsequent input from the level below is low, there is less need for updating the model which would be reflected in reduced sensorimotor activity. We argue such a predictive mechanism can explain the greater mu suppression associated with observation of ambiguous facial expressions. From this perspective, the prediction error is determined by the reciprocal signalling that takes place between the kinematics level in which the movement of the parts of the observed face is described, and the goal level that describes the actor’s short-term goal of making an emotional facial expression. In real life, while decoding facial expressions, we take into consideration social information received from multiple sources, such as bodily gestures, prosody and content of speech. Such contextual information enables us to narrow down the otherwise excessive number of possible emotions that can be observed in a face. In the present study, the two-alternative forced-choice nature of the task reduced the need for contextual information. Participants searched for either of the two emotions (i.e., happiness or sadness). In this scenario, given that the actor’s inferred goal is either to make a happy or a sad face, the observer predicts the associated kinematics on the basis of their own action system: lifting of the lip corners in the former, corrugation of the eyebrows in the latter. The predicted kinematics then get compared to the observed kinematics to produce a prediction error. Based on this prediction error, the observer updates their representation of the actor’s goal, and reaches a decision: “They are making a happy (or sad) face.” The greater discrepancy between the predicted kinematics of the face depicting happiness/sadness and the observed kinematics of the expression in the ambiguous condition leads to more intense sensorimotor activity, indexed as increased mu suppression.

Selective sensitivity of the mu rhythm to the ambiguity of emotion information may be due to the heightened social importance a stimulus gains during focus on its emotional content in order to categorise it as happy or sad compared to, for example, focusing on its direction to categorise it as right- or left-oriented. According to the social-relevance hypothesis of the MNN (Kilner et al., 2006), the predictive MNN is activated only when the observed information is deemed socially relevant (Kilner et al., 2006; Menoret et al., 2015). For example, Kilner et al. (2006) reported greater mu suppression during observation of arm movements when the actor was facing towards the participant rather than facing away from them. They interpreted this result as indicating the role of visuospatial attention as a gating mechanism for the mirroring system, leaving only the socially-relevant information for further processing. Similarly, upon observing a more dynamic temporal pattern in mu rhythm modulation during observation of grasping hand actions in a social context and communicative hand gestures compared to simple grasping actions and meaningless hand gestures, Streltsova et al. (2010) concluded that motor resonance is modulated by the goal and relevance of an action. In line with the social relevance account of the MNN, in the current study, high social significance of emotion information may have activated the sensorimotor cortex differentially in response to easy-to-recognise versus more ambiguous facial expressions. From this perspective, the mirroring system is likely to be engaged during emotion processing, because emotional information is crucial for social interaction. Due to the subjectivity of social relevance, the MNN would not be activated if the social information were perceived to be negligible. Such individual differences in social information processing may partly explain lack of mu suppression observed in some participants in some conditions.

It should be noted that the current EEG results are not able to provide direct evidence for whether the sensorimotor activity plays a causal role in recognising subtle emotional expressions or just reflects this process. Recent findings from a transcranial magnetic stimulation (TMS) study by Paracampo and colleagues (2017) indicate that the sensorimotor area is causally essential for inferring amusement authenticity from observed smiles. Similarly, sensorimotor processing may be necessary for emotion recognition. Brain perturbation techniques could be administered in order to establish such a link between these brain regions and emotion recognition.

### 4.2 Limitations

A major limitation of the study is the small sample size. Due to the limited power of the study, Type I error rate of .05 was assumed per comparison, without any adjustments for multiple tests. Therefore, generalisation of findings is limited and depends on the replication of the present results in future studies with large samples.

### 4.3 Conclusion

We observed increased mu suppression for ambiguous, neutral faces compared to unambiguous happy and sad faces. These results suggest that observing facial expressions may activate a mental simulation network to a greater extent when their emotional content is more ambiguous and difficult to assess. Our findings contribute to the sensorimotor theories of social affective cognition by providing support for the idea that mu suppression during processing of emotional stimuli reflects higher-order cognitive processing associated with emotion understanding, rather than being simply resulting from the observed movement kinematics.

## Author contributions

OK conceived the experiment, performed data acquisition, analysis, interpretation of the data, and wrote and revised the manuscripts. MM contributed substantially to the design and data analysis. NM contributed substantially to the critical revision of the manuscript. IK supervised the study and contributed to the design of the experiment. All authors approved the final draft of the manuscript for publication (OK, MM, NM, IK).

